# Strain identity effects contribute more to *Pseudomonas* community functioning than strain interactions

**DOI:** 10.1101/2024.06.07.597923

**Authors:** Jos Kramer, Simon Maréchal, Alexandre R.T. Figueiredo, Rolf Kümmerli

## Abstract

Microbial communities can shape key ecological services, but the determinants of their functioning often remain little understood. While traditional research predominantly focuses on effects related to species identity (community composition and species richness), recent work increasingly explores the impact of species interactions on community functioning. Here, we conducted experiments with replicated small communities of fluorescent *Pseudomonas* bacteria to quantify the relative importance of strain identity versus interaction effects on two important functions, community productivity and siderophore production. By combining supernatant and competition assays with an established linear model method, we show that both factors have significant effects on functioning, but identity effects generally outweigh strain interaction effects. These results hold irrespective of whether strains interactions are inferred statistically or approximated experimentally. Our results have implications for microbiome engineering, as the success of approaches aiming to induce beneficial (probiotic) strain interactions will be sensitive to strain identity effects in many communities.

## INTRODUCTION

Microbial communities are present in almost all natural and anthropogenic environments on earth, and shape important ecological services from primary production to the decomposition of organic matter and the fixation of greenhouse gases (Stockmann *et al*. 2013; Whitman *et al*. 1998). Given that many of these services can be harnessed for biotechnological applications such as bioremediation and biofuel production (Alper & Stephanopoulos 2009; Minty *et al*. 2013; Piccardi *et al*. 2019; Wang & Chu 2016), recent years have seen a massive surge in studies investigating how contributions of specific microbial players and their interactions with one another affect different functions (collective properties) of microbial communities (Bell *et al*. 2005; Figueiredo *et al*. 2022; Jones *et al*. 2021; Li *et al*. 2019; Ratzke *et al*. 2020; Venail & Vives 2013). While this research has elucidated independent effects of various factors on functions from productivity to invasion resistance, understanding their combined influence is crucial to predicting and manipulating how microbial communities work (Gorter *et al*. 2020; Konopka 2009).

Traditionally, studies on factors shaping community functioning mainly focus on the role of community composition and species richness (Bell *et al*. 2005; Loreau & Hector 2001; Smith & Knapp 2003; Wardle *et al*. 1998). Community composition typically matters because some species may contribute more to functioning than others (Bell *et al*. 2005; Wardle *et al*. 1998). Conversely, species richness typically matters because higher species numbers increase the likelihood of including species with different niche requirements or strong effects on functioning (Bell *et al*. 2005; Loreau & Hector 2001). In both cases, effects on community functioning are ultimately driven by characteristics inherent to specific community members (identity effects). More recently, research has increasingly focused on effects on functioning driven by interactions among community members (interaction effects; Fiegna *et al*. 2015; Figueiredo *et al*. 2022; Gorter *et al*. 2020; Gu *et al*. 2020b; Li *et al*. 2019, 2022; Ratzke *et al*. 2020). In microbes, these interactions are often mediated by secreted products – such as toxins, antibiotic-degrading enzymes, and iron-scavenging siderophores – and can have strong effects on community dynamics (Granato *et al*. 2019; Kramer *et al*. 2020b; West *et al*. 2007). While such interactions are clearly relevant, it remains unclear how their impact on functioning compares to the impact of identity effects because studies typically focus on only one of the two factors (Bell *et al*. 2009; Connolly *et al*. 2013).

Here, we tackle this knowledge gap by quantifying and comparing the impact of strain identity and strain interactions on the functioning of small communities of fluorescent pseudomonads. *Pseudomonas* is a diverse and widespread genus of γ-proteobacteria, occurring in soil and freshwater ecosystems as well as in animal hosts (Cornelis 2010). Fluorescent pseudomonads produce and interact through a broad range of secreted compounds including proteases, biosurfactants and the eponymous fluorescent siderophore pyoverdine (Butaitė *et al*. 2018; Kramer *et al*. 2020a). This versatility has made representatives such as *P. aeruginosa* and *P. fluorescens* important models for studying microbial interactions early on (Brockhurst *et al*. 2007; Griffin *et al*. 2004; Kümmerli *et al*. 2009; Rainey & Rainey 2003). More recently, fluorescent pseudomonads have increasingly been used to study how social traits affect community functioning, including productivity, plant protection, and invasion resistance (Becker *et al*. 2012; Figueiredo *et al*. 2022; Hodgson *et al*. 2002; Hu *et al*. 2016; Jones *et al*. 2021).

In our study, we used 64 diverse soil and freshwater *Pseudomonas* strains belonging to four phenotype classes to examine how strain identity and strain interactions jointly shape two community functions: productivity and the production of iron-scavenging pyoverdines. Pyoverdines are a diverse group of siderophores, and each specific pyoverdine can either promote the growth of strains possessing matching uptake receptors or inhibit the growth of strains without these receptors (Kümmerli 2023). To quantify the positive and negative interactions through pyoverdines and other secreted compounds, we conducted supernatant feeding assays under different conditions. These assays allowed us to calculate community-level interaction metrics. Next, we grew all strains in monocultures and all combinations of two, three, or all four strains per community, and measured their productivity and pyoverdine production over time. This allowed us to leverage an established linear model method (Bell *et al*. 2009) to statistically disentangle the effects of strain identity and strain interactions, and to compare the numerically derived measure of strain interactions with our experimentally measured interaction metrics. Finally, we used our statistical results to identify strains with strong effects on functioning, and tested whether strain interactions had a consistent impact on functioning across communities. Overall, our results show that although strain interactions can significantly affect the functioning of *Pseudomonas* communities, their effects are generally outweighed by those of strain identity.

## MATERIALS AND METHODS

### Strain selection

We drew strains from an established collection of 315 pseudomonads, isolated from eight soil and eight pond samples (18-20 isolates per sample). Sampling and identification of these strains are described elsewhere (Butaitė *et al*. 2017, 2018). Here, we selected a subset of 64 strains comprising four strains per sample [hereafter: community] based on their production of pyoverdine and exo-proteases. While our experiments contrasted conditions where pyoverdine is important for growth with conditions where it is not (see below), we used protease production as a proxy for multidimensional phenotype differences (Kramer *et al*. 2020a) and thereby managed to obtain a highly diverse set of strains (supplementary material; Figure S1, Table S1). Per community, we chose the most divergent strains within the observed phenotype space, aiming to select (i) one strain producing pyoverdine and proteases, (ii) one strain producing only pyoverdine, (iii) one strain producing only proteases, and (iv) one strain producing neither pyoverdine nor proteases (supplementary methods; Figure S2). Our communities thus each featured two strains producing pyoverdine at high levels (the double and the pyoverdine producer; hereafter producers PVD_PRO_ and PVD) and two strains producing no to little pyoverdine (the protease and the non-producer; hereafter non-producers NON_PRO_ and NON). Hereafter, we use ‘strain type’ to refer to those four phenotype classes and ‘strain ID’ to refer to specific representatives of our 64 strains.

### Growth and siderophore production measurements

We quantified growth and pyoverdine production of all strains under iron-limited and iron-rich conditions. First, we grew precultures in 24-well plates with 1.5ml lysogeny broth (LB) per well under static conditions for 48h. Subsequently, we washed cells in 0.85% NaCl and measured their growth ([OD_600_]; optical density measured at 600 nm) against a 0.85% NaCl blank using an Infinite M200 PRO microplate reader (Tecan, Männedorf, Switzerland). Next, we adjusted precultures to an OD_600_ = 0.4 and inoculated 2 µL of each adjusted culture into 96-well plates containing 200 µL medium per well in fourfold replication. We used two variants of CAA medium (5g casamino acids, 1.18g K_2_HPO_4_·3H_2_O and 0.25g MgSO_4_·7H_2_O per liter), an iron-limited variant supplemented with 25 mM HEPES buffer, 20 mM NaHCO_3_ and 100 µg/mL apo-transferrin (a strong iron-chelator), and an iron-rich variant supplemented with 25 mM HEPES buffer and 40 µM FeCl_3_. After 24h of static incubation at 28°C, we quantified growth [OD_600_] and pyoverdine production ([RFU_pvd_]; relative fluorescence units; excitation|emission at 400|460 nm) after 120s of vigorous shaking using the same microplate reader.

### Supernatant assay

We explored interactions through secreted compounds under iron-limited and iron-rich conditions by exposing each strain to its own supernatant and to each supernatant collected from its community members. We harvested supernatants from cultures grown in the above-described experiment by spinning them through 96-well filter plates with a 3 μm glass fiber/0.2 μm Supor membrane (AcroPrep Advance; Pall Corporation, Port Washington, USA) and then collecting the sterile supernatants in 96-well plates. These plates were sealed and stored at -20°C until further use. Next, we grew another set of precultures, washed and adjusted them as before, and subjected them to three treatments: (i) SN_limited_: 180 μL of iron-limited CAA supplemented with 20 μL of supernatant generated under iron-limited conditions; (ii) SN_rich_: 180 μL of iron-rich CAA supplemented with 20 μL of supernatant generated under iron-rich conditions; and (iii) SN_control_: 180 μL of iron-limited or iron-rich CAA supplemented with 20 μL of 0.85% NaCl (mimicking spent medium). Strains were grown in threefold (SN_control_) or fourfold (SN_limited_ and SN_rich_) replication. We measured growth [OD_600_] and pyoverdine production [RFU_pvd_] of each replicate after 24h and 48h of incubation at 28°C under static conditions. We calculated the effects of each supernatant on the producer and each of its community members as growth effects: GE_treatment_ = (SN_treatment_/SN_control_), where SN_treatment_ = SN_limited_ or SN_rich_, with growth values being calculated as the median growth across replicates. Values smaller and greater than one indicate growth inhibition and stimulation, respectively.

We calculated three summary measures of supernatant-based interactions to link supernatant effects to community functioning in direct interactions of multiple strains. Specifically, we calculated for each combination of two, three or all four strains per community, (i) the mean absolute effect and (ii) the proportion of positive effects, thereby separately capturing the strength and sign of supernatant effects, respectively. Additionally, we calculated (iii) an ‘interaction score’ to incorporate information on both strength and sign. To this end, we first tested for each donor-receiver pair whether their reciprocal supernatant effects were positive, negative, or neutral (i.e., whether SN_treatment_ values differed from SN_control_ values; Table S2). Subsequently, we categorized all pairwise interactions based on the effects that the strains had on each other, resulting in six interaction types: mutual stimulation [+/+], one-way stimulation [+/0], no effect [0/0], contrasting effects [+/-], one-way inhibition [0/-], and mutual inhibition [-/-]. Finally, we calculated interaction scores by valuing all effects on other strains [stimulation = 1; neutral effect = 0; inhibition = -1] and then calculating an average score across all interactions for each combination of two, three, or all four strains per community. Interaction scores smaller and greater than zero indicate that inhibitory and stimulatory interactions prevail, respectively.

### Competitions

To be able to assess the effects of strain interactions and strain identity on community functioning, we competed each strain against combinations of its community members under iron-limited and iron-rich conditions, and quantified community productivity and total pyoverdine production over time. Specifically, we set up competition experiments for each of our 16 communities involving all combinations of two, three or all four community members, and included monocultures as controls (15 treatment conditions per community: 4x1 strain + 6x2 strains + 4x3 strains + 1x4 strains). We grew precultures from freezer stocks in 50 ml Falcon tubes containing 5mL LB at 28°C under shaking conditions (170 rpm). After 48h of incubation, we washed cells in 0.85% NaCl, measured the OD_600_ of each culture against a 0.85% NaCl blank, and then adjusted strains to OD_600_ = 0.2. Next, we assembled the mixes and inoculated them at a starting density of OD_600_ = 0.01 either in 6-fold (4-strain competitions) or 5-fold (other treatment conditions) replication into 96-well plates containing 190 µL of iron-limited or iron-rich medium per well. We used a substitutive design, whereby overall starting density is constant across different mixes, while individual strain density decreases when strain number increases (Figueiredo *et al*. 2022; O’Brien *et al*. 2023). We incubated plates in a plate reader at 28°C under static conditions and measured the productivity [OD_600_] and total pyoverdine production [RFU_pvd_] of each culture every 15min over 48h. We used these measurements to calculate integrals of productivity and total pyoverdine production as our primary measures of community functioning. For some analyses, we additionally calculated deviations from expected productivity and pyoverdine production as DEV_trait_ = TV_mix_ – mean(TV_mono_), where TV_mono_ = trait values of monocultures of strains in the mix. The DEV_trait_ values indicate whether the trait value (productivity or pyoverdine production) of a specific strain mix was lower or higher than expected based on the trait values of the monocultures of the constituent strains.

### Linear model method

To compare the effects of strain identity and strain interactions on functioning at the community level, we used an established linear model (LM) method that partitions the variance in a community-level trait between different factors of interest (Bell *et al*. 2009). Briefly, this method uses a series of three LMs to sequentially account for (i) the influence of strain number (entered as a continuous variable), (ii) strain identity (entered as the presence [categorical] of each strain in a particular strain combination), and (iii) strain interactions (strain number, entered as categorical variable). While entering strain number as continuous variable accounts for a linear increase of functioning with strain number, entering it subsequently as categorical variable tests for additional, nonlinear (non-additive) effects, thereby providing a measure of strain interactions among strains (Bell *et al*. 2009). The first LM always uses the focal trait of interest as a response, whereas subsequent LMs are fitted on the residuals extracted from the respective previous model. Note that the effects obtained for strain identity and strain interactions are orthogonal and thus independent of the order in which the corresponding LMs are fitted (Bell *et al*. 2009). We ran LMs separately for each community and experimental condition on data from all 11 combinations of two or more strains, focusing on community productivity and community pyoverdine production as our primary traits of interest. When examining the contributions of different strain types to community functioning, we included DEV_productivity_ and DEV_pyoverdine_ as additional traits to identify strain types driving positive or negative deviations of functioning from its expected value.

To examine the relative importance of strain interactions and strain identity, we extracted the mean squares from the relevant LMs [ii + iii] (Bell *et al*. 2009). Given that the non-linear richness term provides a purely statistical proxy of strain interactions, we additionally considered our summary measures of supernatant-based interactions. To this end, we replaced the strain richness term in the last LMs [iii] by, respectively, the interaction score or the mean absolute supernatant effect as well as the ratio of positive supernatant effects, then extracted the corresponding mean squares, and finally included them together with the mean squares for strain identity and non-linear strain richness in an across-community comparison (see below). To be able to examine whether specific strain types contributed disproportionally to community functioning, we extracted the linear model coefficients obtained for each strain from the LMs [ii] focusing on strain identity. These coefficients provide a measure of each strain’s effect on functioning relative to that of the average strain in the community (Bell *et al*. 2009).

### Statistical analysis

To examine growth profiles and supernatant-based interactions, we tested whether growth or pyoverdine production differed between strain types (PVD_PRO_, PVD, NON_PRO_, or NON) and conditions (iron-rich or iron-limited). Next, we tested whether supernatant effects (GE_treatment_) differed between conditions or supernatant donor and receiver types. To examine the relative importance of strain interactions and strain identity on productivity and pyoverdine production, we tested for differences between the mean square values obtained from our linear model decomposition for strain identity, non-linear strain richness, the interaction score, and the combination of mean absolute supernatant effect and the ratio of positive effects (see above), accounting for differences between conditions. Similarly, we examined the contributions of different strain types to (deviations from expected) community functioning by comparing the strain coefficients obtained through the linear model method across our communities. To examine whether strain interactions consistently shape functioning across communities, we finally tested whether deviations from expected productivity (DEV_productivity_) were shaped by strain number (2, 3, or 4; categorical), condition, or the interaction score summarizing supernatant-based interactions in the different sets of strains. Note that we always included the strains’ habitat-of-origin (soil or pond) as a co-factor into our models, but do not report the corresponding results here because they do not affect our main results and were rarely significant (all results are reported in the supplementary material).

We implemented our analyses in R 4.2.1 (www.r-project.org) using LMs, generalized least squares (GLS) models and linear mixed models (LMMs). GLS models and LMMs were implemented using the *gls* and *lme* functions (nlme package; Pinheiro et al., 2023). We obtained p-values of effects in these models using the *Anova* function (car package; Fox & Weisberg, 2019). We used the emmeans package (Lenth, 2021) to perform post hoc analyses and adjusted p-values for multiple testing (n_test_ > 2) using the false discovery rate. Unless otherwise stated, models were initially fitted with all possible interaction terms. Where required, we transformed response variables to obtain normally distributed residuals. To account for the non-independence of strains from the same community and for multiple measurements of each strain under different conditions, we initially fitted all models as random intercept models using community and, in case of repeated measurements, strain (mixture) identity nested within community as random effect(s). Each final model was then selected in a two-step procedure. First, we used the Akaike information criterion (AIC) to simplify the random effect structure and to select an appropriate variance structure (using the weights-argument in the *gls* and *lme* function) where residual plots indicated a deviation from homogeneity (Zuur *et al*. 2009). Second, we simplified the fixed component by dropping non-significant interaction terms (p > 0.05). The structure of all final models is detailed in Table S3.

## RESULTS

### Growth and pyoverdine production profiles of *Pseudomonas* strains

We first confirmed that pyoverdine producers (PVD_PRO_ and PVD) and non-producers (NON_PRO_ and NON) behaved as expected by quantifying pyoverdine production and growth under iron-limited and iron-rich conditions. Indeed, we found that the producer types featured higher pyoverdine production than the non-producer types and that pyoverdine favors growth when iron is scarce (Table 1+S4, Figure 1). Specifically, PVD_PRO_ and PVD grew better than the non-producers under iron limitation, while NON_PRO_ reached higher densities than NON. Somewhat surprisingly, the growth differences between PVD and NON_PRO_ were small despite marked differences in pyoverdine production (Figure 1A+B), suggesting that NON_PRO_ might produce secondary siderophores such as pyochelin (Kümmerli 2023). Indeed, when measuring the total iron-chelating activity using the cholorimetric CAS assay (Schwyn & Neilands 1987), we found that NON_PRO_ featured the same activity as PVD (supplementary material; Table S5, Figure S3). Under iron-rich conditions, where siderophores are not required for sustained growth, all strain types reached similar densities (Table 1A, Figure 1A) and produced little pyoverdine (iron-rich vs. iron-limited: PVD_PRO_: t_70_ = -14.20, p < 0.001; PVD: t_70_ = -12.98, p < 0.001; NON_PRO_: t_70_ = -4.35, p < 0.001; NON: t_70_ = -1.19, p = 0.239, Figure 1B).

**Figure 1.**
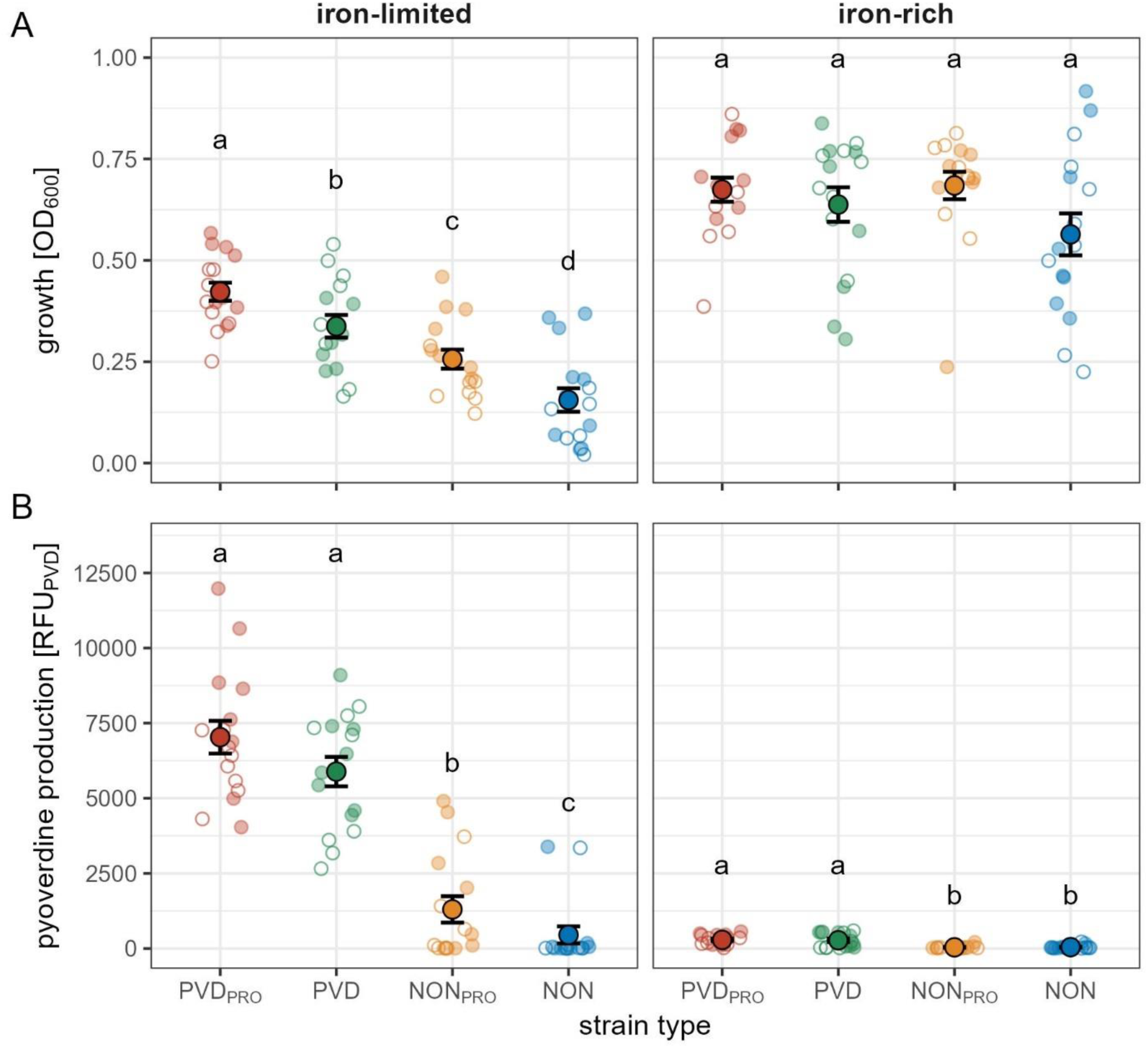
Strain types differ in their growth and pyoverdine production profiles. (A) Growth and (B) pyoverdine production of PVD_PRO_ (red), PVD (green), NON_PRO_ (orange), and NON (blue) strains isolated from eight soil (empty small circles) and eight freshwater (filled small circles) communities (one strain per type and community), measured in iron-limited and iron-rich medium. Small circles represent the median of four replicates obtained for each strain under each condition. Large circles and black lines show mean and standard error, respectively. Letters show significantly different types. All comparisons were performed within each medium (detailed statistical results are provided in Table S4).

**Table 1.** Growth and siderophore production profiles. Determinants of (A) monoculture growth and (B) pyoverdine production of soil and freshwater *Pseudomonas* strains belonging to four different strain types varying in their production of proteases and the siderophore pyoverdine (PVD_PRO_, PVD, NON_PRO_, and NON).

|  | (A) growth |  |  | (B) pyoverdine production |  |  |
| --- | --- | --- | --- | --- | --- | --- |
| | df | $\chi^2$ | p | df | $\chi^2$ | p |
| habitat | 1 | 5.28 | **0.022** | 1 | 3.01 | 0.083 |
| medium | 1 | 217.93 | **< 0.001** | 1 | 267.68 | **< 0.001** |
| type | 3 | 65.90 | **< 0.001** | 3 | 115.62 | **< 0.001** |
| habitat : type | 3 | 9.75 | **0.021** | - | - | - |
| medium : type | 3 | 9.79 | **0.021** | 3 | 122.89 | **< 0.001** |

### Supernatant effects are pronounced under iron-limited conditions

Next, we harvested supernatants from each strain and measured their effects on the growth of community members and the producers themselves (Table S6+S7, Figure 2). We first focus on effects that the supernatants had on their own producers. Under iron-limited conditions, ‘effects on self’ were pronounced and overall positive for all types, indicating that strains typically secrete pyoverdine or other compounds into the supernatant that favor their own growth. This was not the case under iron-rich conditions, where effects on self were typically neutral or negative and generally small (Table S7, Figure 2). Focusing next on the impact of supernatants on other community members, we likewise found that effects under iron-rich conditions were neutral or negative and typically small, suggesting a low baseline production of toxic compounds. Under iron-limited conditions, however, supernatant effects on others varied substantially, ranging from strong inhibition to strong stimulation for specific strain combinations (Table S7, Figure 2). To further examine how interactions differ between iron conditions, we classified all pairwise interactions within a community, ranging from mutual inhibition to mutual stimulation, and displayed them as individual interaction heatmaps for each community and condition (Figure 3). This qualitative analysis revealed that there are many more positive interactions among strains under iron-limited than iron-rich conditions.

**Figure 2.**
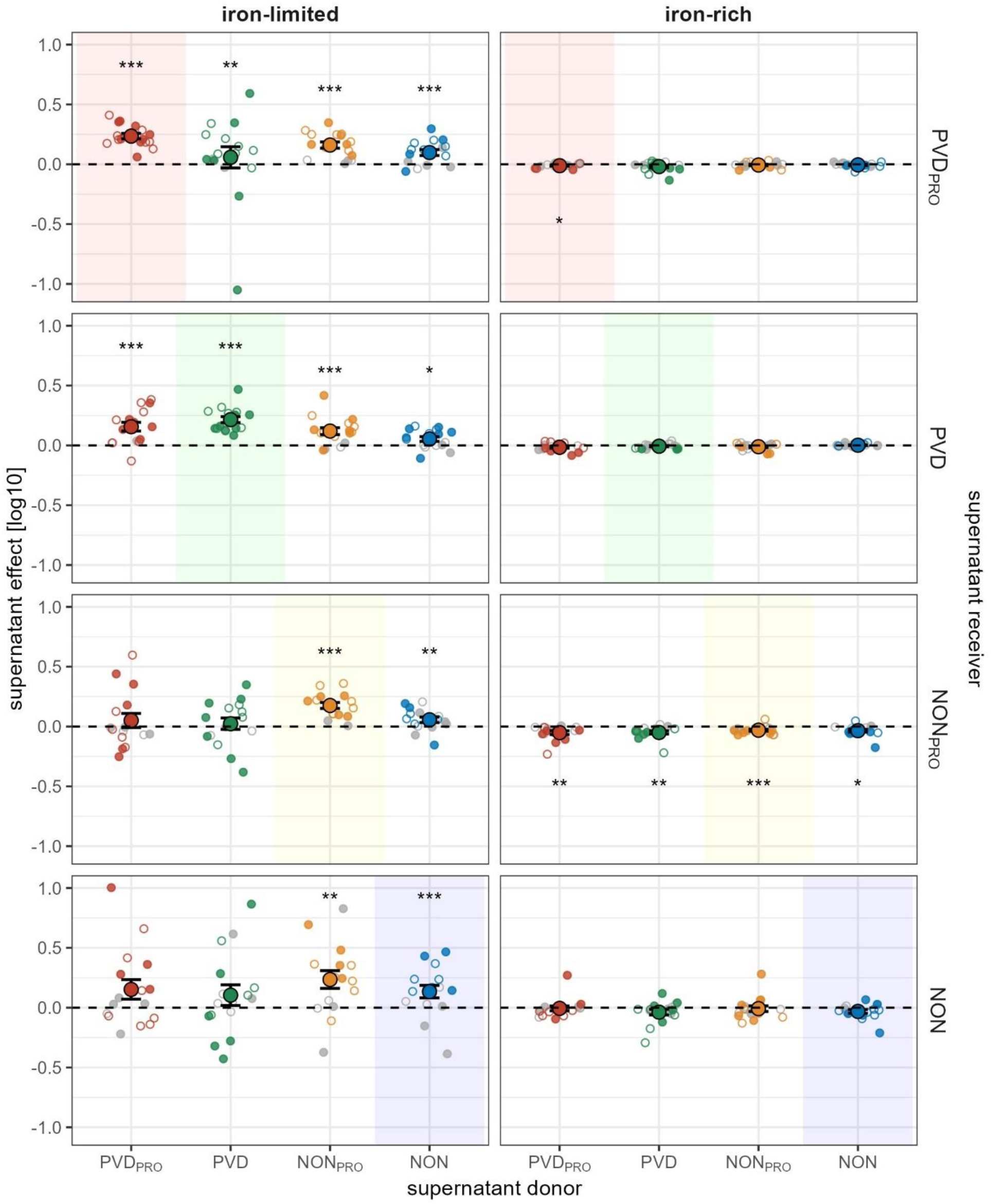
Secreted compounds have pronounced effects under iron-limitation. Shown are effects that PVD_PRO_, PVD, NON_PRO_, and NON strains isolated from soil (empty small circles) and pond (filled small circles) communities have on each other’s growth through compounds secreted into the supernatant under iron-limited and iron-rich conditions. Small circles show the median of four replicates obtained for each donor/receiver combination. Small grey circles show supernatant effects of specific donor/receiver combinations that did not differ from neutrality, whereas small colored circles indicate significant effects on receiver growth. Large circles and black lines show mean and standard error, respectively. Dashed horizontal lines indicate the null line where compounds in the supernatant have no effect on receiver growth. Colored rectangles highlight the effects that strains have on their own growth. Asterisks above and below the null line indicate that the average supernatant effect of a specific combination of donor and receiver types was significantly positive and negative, respectively (significance levels are indicated as follows: * 0.05 ≥ p > 0.01; ** 0.01 ≥ p > 0.001; *** p ≤ 0.001; see Table S6+S7 for further details).

**Figure 3.**
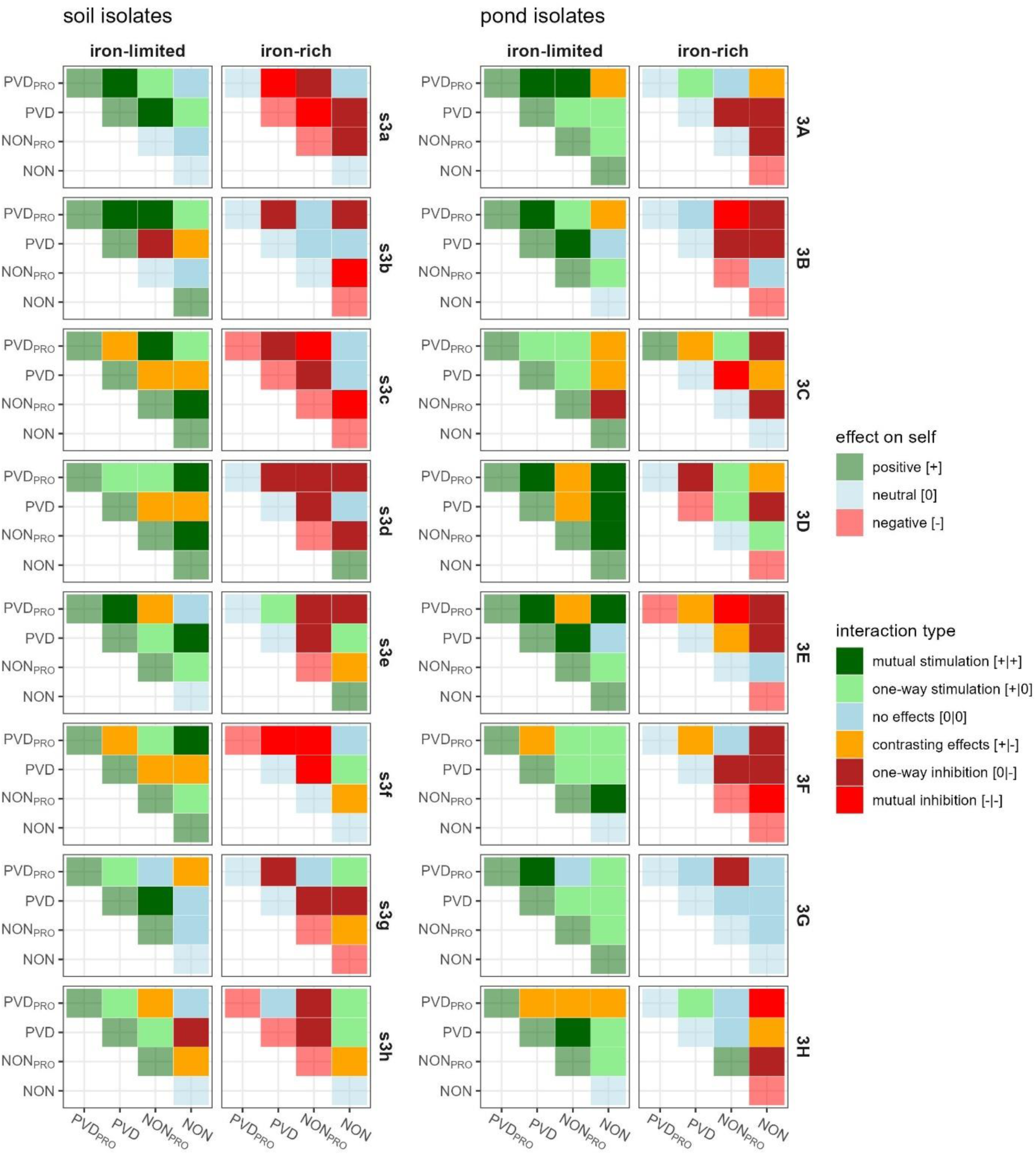
Interaction types illustrate the high potential for growth stimulation under iron limitation. Interaction types between – and effects on self of – PVD_PRO_, PVD, NON_PRO_, and NON strains from eight soil (s3a to s3h) and eight freshwater (3A to 3H) communities of *Pseudomonas* bacteria. Interaction types (opaque colors) and effects on self (transparent colors) were assigned based on the positive, neutral, or negative effects that the strains had on each other and themselves through compounds secreted into the supernatant under iron-limited and iron-rich conditions (see Figure 2 and the Methods for details).

### Identity effects explain more variation in community functioning than strain interactions

We examined the effects of strain identity and interactions on two metrics of community functioning, productivity and pyoverdine production. We set up monocultures and all combinations of two, three, or all four strains per community and used an established linear model method (Bell *et al*. 2009) to determine the variance explained by strain identity and strain interactions. We considered three measures of strain interactions: (i) non-linear strain richness, a statistical proxy for interactions (Bell *et al*. 2009); (ii) the interaction score, a proxy reflecting the overall sign of supernatant effects within communities (Figure 3); and (iii) a combination of the mean absolute supernatant effect and the proportion of positive effects, which accounts for both magnitude and sign of strain interactions.

We found that strain identity explained more variation in community productivity (Figure 4A) and pyoverdine production (Figure 4B) than strain interactions, regardless of which measure of strain interactions we considered, and independent of the iron condition (Table 2+S8). When comparing the three measures of strain interactions, we found that the combination of mean absolute supernatant effect and the proportion of positive supernatant effects explained more variation in functioning than both non-linear strain richness and the interaction score (Table 2, Figure 4). Independent of these effects, strain identity explained more variation in productivity – and all predictors explained more variation in pyoverdine production – under iron-limited than iron-rich conditions (Table S8, Figure 4).

**Figure 4.**
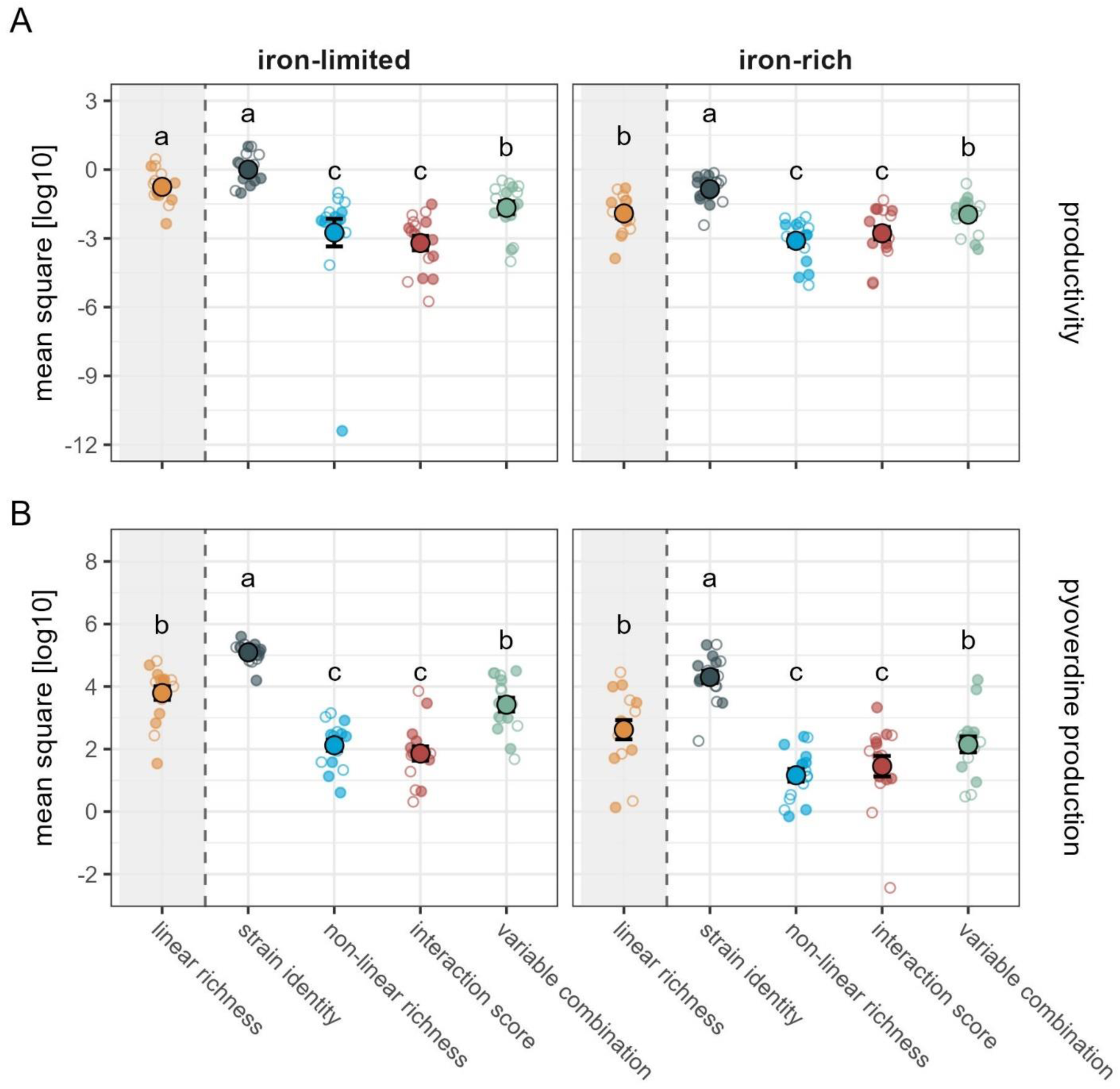
Strain identity explains more variation in community productivity than strain interactions. Shown are mean square values extracted from linear models fit for each of eight soil (empty small circles) and eight pond (filled small circles) communities to explain the impact of strain identity and three measures of strain interactions, non-linear richness, the interaction score, and a combination of variables on (A) community productivity and (B) pyoverdine production. The impact of linear richness, i.e. the extent to which community functioning linearly increases with strain number, is shown for comparison (grey-shaded area). While non-linear richness is a purely statistical proxy for strain interactions, the interaction score and the combination of variables are based on supernatant effects. The interaction score predominantly reflects the sign of supernatant effects, whereas the combination of variables includes the mean absolute supernatant effect and the proportion of positive supernatant effects and thus separately accounts for both sign and magnitude. Large circles and black lines show means and standard errors. Small circles show mean square values obtained for specific communities from models fit separately to data generated under iron-limited and iron-rich conditions (letters show significantly different impacts on functioning based on the results of our statistical models; see the Methods and Table 2+S8 for details).

**Table 2.**
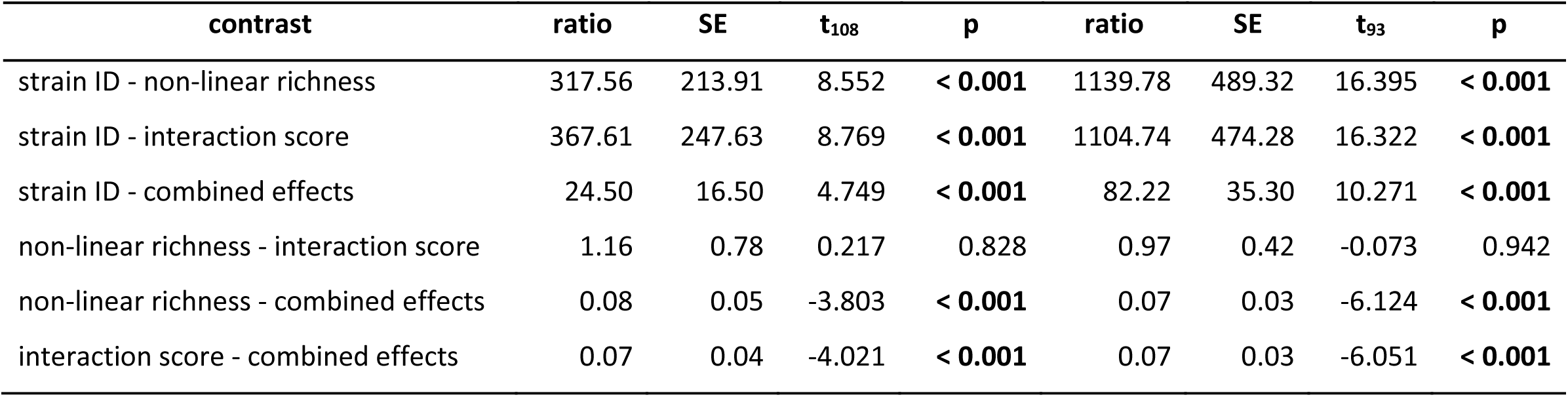
Impact of strain identity and interactions on community functioning. Post-hoc comparisons of different determinants of (A) productivity and (B) pyoverdine production of 16 small *Pseudomonas* communities. Significant p-values are in bold.

| contrast | (A) productivity |  |  |  | (B) pyoverdine production |  |  |  |
| --- | --- | --- | --- | --- | --- | --- | --- | --- |
|  | ratio | SE | t<sub>108</sub> | p | ratio | SE | t<sub>93</sub> | p |
| strain ID - non-linear richness | 317.56 | 213.91 | 8.552 | < **0.001** | 1139.78 | 489.32 | 16.395 | < **0.001** |
| strain ID - interaction score | 367.61 | 247.63 | 8.769 | < **0.001** | 1104.74 | 474.28 | 16.322 | < **0.001** |
| strain ID - combined effects | 24.50 | 16.50 | 4.749 | < **0.001** | 82.22 | 35.30 | 10.271 | < **0.001** |
| non-linear richness - interaction score | 1.16 | 0.78 | 0.217 | 0.828 | 0.97 | 0.42 | -0.073 | 0.942 |
| non-linear richness - combined effects | 0.08 | 0.05 | -3.803 | < **0.001** | 0.07 | 0.03 | -6.124 | < **0.001** |
| interaction score - combined effects | 0.07 | 0.04 | -4.021 | < **0.001** | 0.07 | 0.03 | -6.051 | < **0.001** |

### Strain types vary in their impact on community productivity

Above, we have shown that strain identity effects outweigh the effects of strain interactions on community functioning. However, this result does not reveal whether strain types differ in their impact on functioning. To tackle this question, we again leveraged the above-described linear model method, this time focusing on the ‘strain ID’ coefficients extracted from models using either community productivity, pyoverdine production, or the deviations from their expected values as functions of interest. The strain ID coefficients provide a measure of each strain’s effect on community functioning relative to an average strain’s contribution (Bell *et al*. 2009).

We found that PVD_PRO_ made above-average contributions to community productivity across iron conditions (main effect of type: χ^2^ = 23.18, p < 0.001; PVD : t = 2.67, p = 0.009; Figure 5A), while NON contributed less than the average (t_82.7_ = -3.89, p < 0.001; Figure 5A). Intriguingly, the presence of NON strains was generally associated with higher-than-average deviations from expected productivity (main effect of type: χ^2^ = 7.89, p = 0.048; NON: t_124_ = 2.36, p = 0.020; Figure 5B). When focusing on pyoverdine production, we unsurprisingly observed that pyoverdine producers and non-producers made, respectively, above-average and below-average contributions across iron conditions (main effect of type: χ^2^ = 138.85, p < 0.001; type-condition interaction: χ^2^ = 29.81, p < 0.001; Table S9; Figure 5C). In addition to these strain type effects, we observed that individual strains of each strain type variously featured above-average or below-average contributions to all measures of functioning (Table S10; Figure 5).

**Figure 5.**
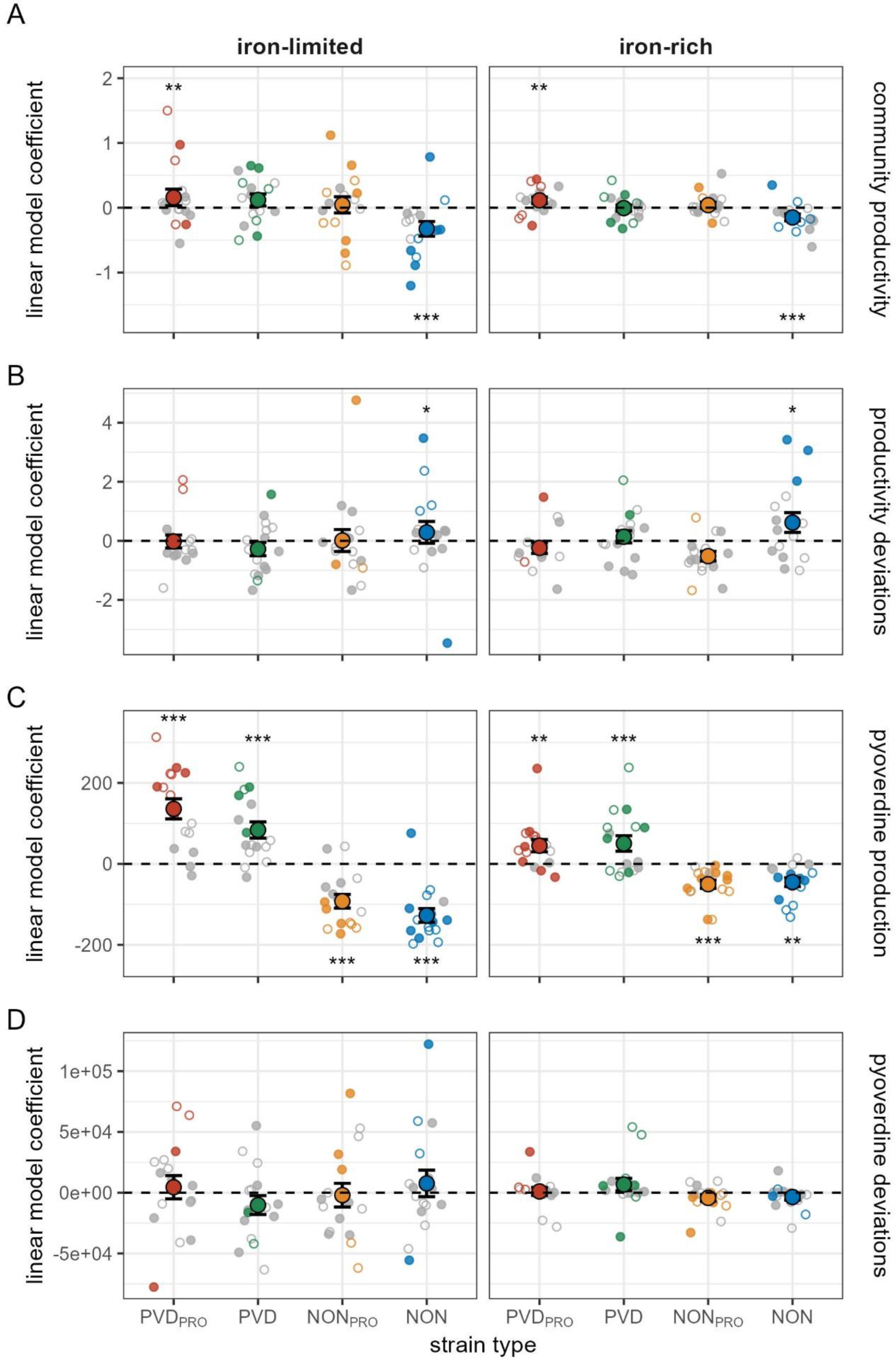
Strain types and individual strains differ in their impact on community productivity. Shown are linear model coefficients indicating the effect that PVD_PRO_ (red), PVD (green), NON_PRO_ (orange), and NON (blue) strains from soil (empty small circles) and pond (filled small circles) communities had, relative to the average in their community, on (A) community productivity, (B) deviations from expected community productivity, (C) pyoverdine production, and (D) deviations from expected pyoverdine production. Large circles and black lines show means and standard errors. Dashed lines indicate average strain effects. Small circles show linear model coefficients obtained for specific strains from models fit separately to data generated for each community under iron-limited and iron-rich conditions, respectively. Small grey circles indicate that linear model coefficients were not significant, whereas small colored circles indicate significant coefficients. Asterisks above and below the average effect line indicate that the average effect of a specific type was significantly positive and negative, respectively (significance levels are based on the results of our statistical models and indicated as follows: * 0.05 ≥ p > 0.01; ** 0.01 ≥ p > 0.001; *** p ≤ 0.001; statistics for specific strains are given in Table S10).

### Interaction scores can predict productivity across communities

Our findings show that strain interactions have a smaller impact on functioning than strain identity, but this does not mean that strain interactions are insignificant. In a last step, we therefore tested whether strain interactions had a consistent impact on functioning across communities. We focused on deviations from expected productivity as function of interest and used the interaction score as a measure of interactions among community members. We predicted that low (inhibitory) and high (stimulatory) interaction scores should be associated with reduced and increased productivity deviations, respectively. In line with this idea, we found a positive relationship between deviations from expected productivity and the interaction score (Table 3; slope ± SE: 0.46 ± 0.18, t_331_ = 2.59, p = 0.010, Figure 6A). Independent of this effect, productivity deviations were higher under iron-rich than under iron-limited conditions (Table 3; soil: t_331_ = 8.13, p < 0.001; pond: t_331_ = 3.91, p < 0.001; Figure 6B) and increased with strain number (Table 3; two vs. three strains: t_331_ = 3.67, p < 0.001; three vs. four strains: t_331_ = 2.36, p = 0.019; Figure 6C).

**Figure 6.**
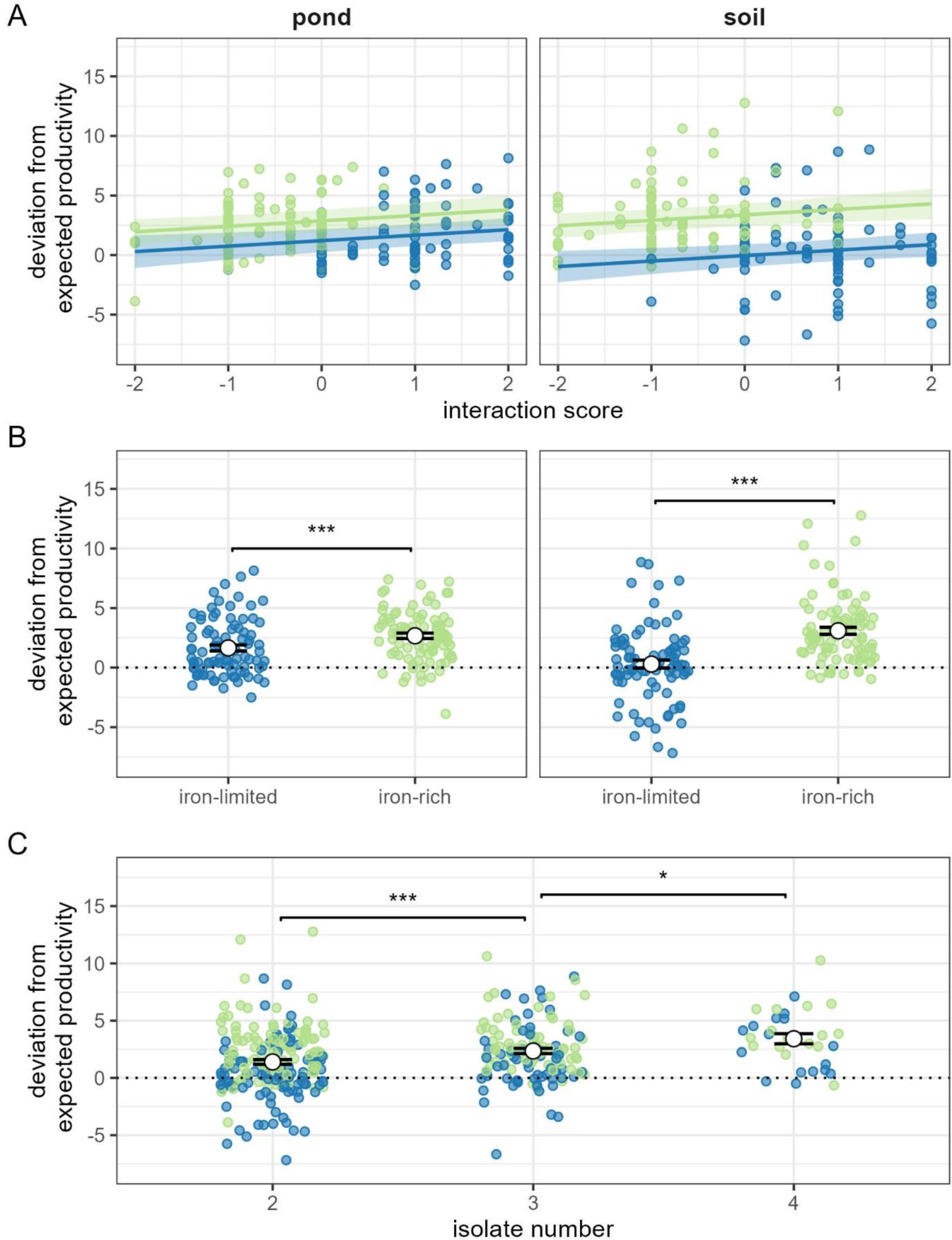
Deviations from expected productivity increase as supernatant-based interactions shift from inhibitory to stimulatory. Shown are relationships between deviations from expected community productivity and (A) the supernatant-based interaction scores, (B) iron condition, and (C) strain number, as measured under iron-limited (blue) and iron-rich (green) conditions in pond and soil strains [left and right panels in (A) and (B), respectively]. High interaction scores indicate that stimulatory effects of secreted compounds prevail, whereas low interaction scores indicate a prevalence of inhibitory effects. Solid lines and shaded areas in (A) are regression lines and 95% confidence intervals, respectively. Large white circles and black lines in (B) and (C) show means and standard errors, respectively. Significance levels in (B) and (C) are indicated as follows: * 0.05 ≥ p > 0.01; ** 0.01 ≥ p > 0.001; *** p ≤ 0.001.

**Table 3.**
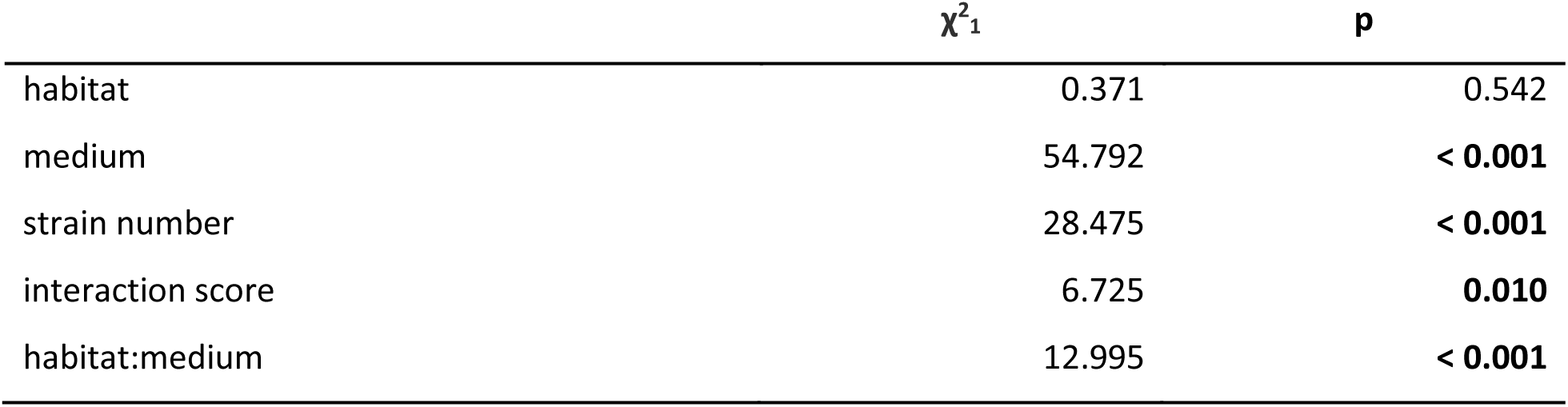
Deviations from expected community productivity. Determinants of deviations from expected community productivity. Significant p-values are in bold.

| | $\chi^2_1$ | p |
| --- | --- | --- |
| habitat | 0.371 | 0.542 |
| medium | 54.792 | **< 0.001** |
| strain number | 28.475 | **< 0.001** |
| interaction score | 6.725 | **0.010** |
| habitat:medium | 12.995 | **< 0.001** |

## Discussion

Although microbial communities shape critical ecosystem services, such as primary production and the fixation of greenhouse gases, we are only beginning to understand how these communities function. Here, we examined and compared the impact of strain interactions and strain identity on the functioning of 16 communities of *Pseudomonas* bacteria, each comprising four strains belonging to four distinct types varying in their potential to secret the siderophore pyoverdine and proteases. We found that strain identity effects explained more variation in community productivity and pyoverdine production than strain interactions, regardless of whether interactions were inferred statistically or measured more directly using supernatant assays. While different strain types overall affected functioning in accordance with their baseline (monoculture) growth and pyoverdine production, we also found that individual strains deviated from this pattern in co-culture, suggesting that strain interactions can modulate the impact of strain identity. In line with this idea, we found that deviations from expected productivity consistently increased across communities as interactions through secreted compounds shifted from inhibitory to stimulatory. Altogether, our findings suggest that although effects of strain identity may often outweigh the effects of strain interactions, both factors will usually be required to gain a nuanced understanding of how naturally diverse communities function.

Classical studies of sociomicrobiology focusing on closely related strains have consistently reported that effects of strain interactions can outweigh the impact of strain identity on functioning (e.g., (Gore *et al*. 2009; Griffin *et al*. 2004; Sandoz *et al*. 2007; Strassmann *et al*. 2000). For instance, (Griffin *et al*. 2004) showed that a pyoverdine-producing *P. aeruginosa* strain grew to higher density than a closely related non-producer in monoculture, but was exploited and eventually outcompeted by that non-producer in mixed culture. By contrast, we found that interactions between genetically more diverse natural *Pseudomonas* strains have comparably smaller effects on functioning and are typically outweighed by pronounced effects of strain identity. Identity effects often result from differences in individual-level traits such as growth rate and metabolic capabilities (Lajoie & Kembel 2019) and such differences are expected to increase as strains become genetically less similar (Bernhardsson *et al*. 2011; Oña *et al*. 2021). Our findings thus suggest that the impact of strain interactions relative to strain identity declines when moving from genetically homogenous to more diverse communities characterized by lower relatedness.

We have previously shown that the siderophores expressed by *Pseudomonas* isolates mediate diverse and strong strain interactions (Figueiredo *et al*. 2022), and our supernatant assays indeed revealed a greater scope for such interactions under iron-limited than iron-rich conditions. We therefore initially expected strain interactions to have a higher impact on community functioning under iron limitation. Contrary to this expectation, we found little evidence that different iron conditions affected the relative impact of social interactions on community functioning. A potential explanation for this pattern could be social trait linkage. We have previously shown that the production of different secreted compounds, including pyoverdines, proteases, biofilm and toxic compounds, are often positively correlated among our natural isolates (Kramer *et al*. 2020a). This would mean that strains contributing a lot to pyoverdine production under iron limitation may contribute a lot to other secreted compounds under iron rich conditions, leading to consistent (large or small) effects of specific strains on functioning across conditions. In support of this idea, we found that PVD_PRO_ strains contributed more – and NON strains less – to productivity than the average strain regardless of iron condition. Hence, positive trait linkage may stabilize the impact of individual community members on functioning across environmental conditions.

We found that deviations from expected productivity increased when supernatant effects shifted from inhibition to stimulation, indicating that mutually stimulatory effects of secreted compounds could promote community functioning. One possible explanation is that such mutual stimulation entails mutual benefits, which can occur if it increases the availability of a growth-limiting resource for strains with non-overlapping niche requirements (Finke & Snyder 2008; Oña *et al*. 2021). However, mutual stimulation often results in mutual exploitation between competing strains, where each strain strives to increase its own fitness at the expense of the other by using its secreted, publicly available compounds without paying the associated costs (Oliveira *et al*. 2014). Such mutual exploitation (or cheating; (Ghoul *et al*. 2014)) could promote higher than expected functioning at the community level if one strain derives net benefits from the interaction that outweigh the net costs to the other strain(s) (MacLean *et al*. 2010; Mridha & Kümmerli 2022). Overall, these considerations highlight that the secretion of sharable ‘public goods’ might often strongly affect the functioning of bacterial communities even when identity effects are strong.

To quantify the effect of strain interactions on community functioning, we used non-linear strain richness as a statistical proxy for strain interactions (Bell *et al*. 2009) and compared its impact with that of more direct interaction measures based on supernatant effects. We found that non-linear richness performed as well as our univariate interaction score, but captured less variation in functioning than a combination of two variables separately capturing the magnitude and sign of supernatant effects. These findings validate both non-linear richness and the interaction score as simple measures of strain interactions. Moreover, they show that experimental data can provide additional benefits in estimating the impact of strain interactions on functioning, but only if the strength and direction of interactions are separately accounted for. We therefore recommend obtaining experimental data when the impact of strain interactions is of particular interest, but to rely on non-linear richness as proxy when it is not.

In recent years, interest in approaches to manipulate microbial assemblies and microbiomes to perform beneficial functions has skyrocketed (Gu *et al*. 2020a; Ibrahim *et al*. 2021; Inda *et al*. 2019; Mueller & Sachs 2015) and the use of probiotics has become popular in agriculture (Menendez *et al*. 2017), aquaculture (Verschuere *et al*. 2000) and human health (Kerry *et al*. 2018). Many of the proposed approaches make use of microbial interactions with the idea to introduce or promote probiotic strains in communities that engage in resource or interference competition with pathogens (Hu *et al*. 2016) or exploit their social traits (Brown *et al*. 2009). Such interactions are often driven by secreted secondary metabolites such as toxins (interference competition; Hu *et al*. 2016) and siderophores (siderophore exploitation; González *et al*. 2021). While these approaches certainly hold great promise, our results highlight that the introduction of cheating mutants might only be successful if their individual-level characteristics allow the mutants to persist in their niche even when the pathogen they exploit declines in frequency. More generally, our findings suggest that strain identity effects, in addition to social interactions, are key to consider in order to develop powerful and sustainable probiotics.

In conclusion, we showed that strain identity effects have a larger impact on the productivity and pyoverdine production of *Pseudomonas* communities than strain interactions. Our results suggest that strain interactions might often reinforce or diminish, but rarely overwrite, existing baseline differences in how diverse microbial players affect the functioning of their community. While our study solely focused on *Pseudomonas* bacteria, we anticipate even stronger identity effects on our functions of interest in taxonomically more diverse microbial communities. At the same time, we note that the patterns observed here might change for other functions. Productivity often resembles a zero-sum game where one strain can only increase its contribution to functioning at the expense of another. By contrast, other functions may leave more scope for net-positive complementation (e.g., community respiration; Bell *et al*. 2009) and these functions should be determined to a greater extent by species interactions relative to identity effects. Future work should therefore consider multiple functions in addition to productivity to unravel the importance of species interactions in natural microbial communities of varying taxonomical diversity.

### Author contributions

Conceptualization: JK and RK; Experiments: JK, SM and ARTF; Data Analysis: JK and ARTF; Writing (first draft): JK and RK; Writing (refinement): JK, SM, ARTF and RK.

## Acknowledgements

We thank Elena Butaitė for collecting the natural isolates.

