## Supplementary Material for "Strain identity effects contribute more to *Pseudomonas* community functioning than strain interactions"

This file contains the following Supplementary Information:

- **Supplementary Methods**
- **Supplementary Analyses**
- **Supplementary References**
- **Supplementary Table S4-S9**

[Tables S1, S2, S3 and S10 are avialable as separate files]

- **Supplementary Figures S1-S3**

**Supplementary Methods**

**Strain selection.** We selected strains from an established collection of 315 *Pseudomonas* strains isolated from eight soil and eight freshwater samples (18-20 strains per sample; hereafter: community) (Butaitė *et al.* 2017, 2018). From each of these 16 communities, we selected four strains based on their production of pyoverdine and exo-proteases. Per community, we chose the most divergent strains within the observed phenotype space, aiming to select (i) one strain producing pyoverdine and proteases [PVD_PRO_], (ii) one strain producing only pyoverdine [PVD], (iii) one strain producing only proteases [NON_PRO_], and (iv) one strain producing neither pyoverdine nor proteases [NON]. To quantify pyoverdine production, we grew all 315 strains in iron-limited casamino acids medium and measured the natural fluorescence of pyoverdine in the culture supernatant. To quantify protease production, we spotted aliquots of bacterial culture onto skim milk agar and then measured the size of the proteolytic halo forming around protease-producing colonies (see Kramer *et al.* 2020 for further details). Based on the pyoverdine and protease production values, we assigned one of the four types to each strain, and then picked one strain per type for each community. When the available types were not clearly defined (e.g., because no complete non-producers of pyoverdine and proteases where present in the focal set of strains), we picked a strain with a profile close to the intended type. When possible, we included previously sequenced strains (see Butaitė *et al.* 2017). Overall, the pyoverdine/protease production of our selected strains matched the intended type, with PVD_PRO_ and PVD typically producing pyoverdine at much higher levels than NON_PRO_ and NON (Figure S2).

To verify that we selected a diverse set of strains, we quantified the phylogenetic distance between strains of the same habitat (soil or pond) based on partial sequences of the *rpoD* gene. To this end, we first obtained the partial *rpoD* gene sequences of the selected strains from the European Nucleotide Archive (generated by Butaitė *et al.* 2017; Accession number: PRJEB21289). Next, we used the DistanceMatrix function (DECIPHER package; Wright 2016) to calculate the pairwise dissimilarity between, respectively, all pond and all soil isolates, and then transformed the default dissimilarity measures into similarity measures using a “1-Matrix” operation. To assess the diversity of strains within our experimental communities, we checked whether strains from the same community belonged to the same or different species, using a pairwise similarity threshold of 96% to identify conspecifics (Mulet *et al.* 2017; Sánchez *et al.* 2014). We found that strains within communities generally represented different species, with only one exception (in soil community s3d, PVD and NON_PRO_ showed a pairwise similarity of 97% despite marked differences in their phenotype; Table S1, Figure S2).

To display the phylogenetic diversity of our Pseudomonas strains, we generated phylogenetic trees for soil and pond strains (Figure S1). First, we obtained *rpoD* gene sequences of five reference *Pseudomonas* strains from the Pseudomonas.com database (Winsor *et al.* 2016). We then used the subseq function (XVector package; Pagès & Aboyoun 2024) to trimm the full-length *rpoD* sequences of these type-strains at positions 488 and 1044, so that their length would match the partial sequences of our natural isolates. Next, we generated two alignment objects (for pond isolates and soil isolates, each with the type strains) using the AlignSeqs function (DECIPHER package), and then ran the alignments through the phyDat function (phangorn package; Schliep 2011) to transform them into phyDat objects. Finally, we ran the phyDat objects through the modelTest function and used the output to generate Maximum Likelihood trees with the pml_bb function, subsequently bootstraped with the bootstrap.pml function (all phangorn package). After rooting the trees at their midpoints (midpoint function, phangorn package), we generated the tree figures with the ggtree package (Yu *et al.* 2017).

**Quantification of total siderophore production.** To quantify the total production of siderophores per strain (type), we used the Chrome Azurol Sulfonate (CAS) assay (Schwyn & Neilands 1987). This colorimetric assay uses an Fe(III)-Chrome Azurol Sulfonate-Hexadecyltrimethylammonium bromide [Fe(III)-CAS-HDTMA] complex that changes color, from blue to orange, when it is deferrated by exchange reactions with siderophores. We first collected supernatant generated by each strain under iron-limited and iron-rich conditions (details in the main text), and then transferred 20 µL of each supernatant into 240 µL of CAS assay solution (prepared according to the original recipe; see Schwyn & Neilands 1987). We incubated these mixtures at room temperature in the dark for 30 minutes, and then quantified the total iron-chelating activity, a proxy measure of the total production of siderophores, by measuring the loss of the blue-colored Fe-CAS-HDTMA complex at 630 nm.

**Supplementary Analysis**

**Total siderophore production of strain types.** In the main text, we show that pyoverdine production differs between producer (PVD_PRO_ and PVD) and non-producer types (NON_PRO_ and NON). However, pseudomonads sometimes deploy additional siderophores to scavenge for iron. To account for this possibility, we measured the total iron-chelating activity of each strain as a proxy for the production of all siderophores (see above), and then examined whether it differed between types, experimental conditions or the habitat-of-origin. We found that all these factors jointly affected siderophore production (habitat: χ^2^_1_ = 0.63, p = 0.427; condition: χ^2^_1_ = 348.18, p < 0.001; donor type: χ^2^_3_ = 0.06, p = 0.996; interaction habitat x donor type: χ^2^_3_ = 8.72, p = 0.033; interaction condition x donor type: χ^2^_3_ = 49.25, p < 0.001). Under iron-limited conditions, PVD_PRO_ and NON featured, respectively, the highest and lowest siderophore production, while PVD and NON_PRO_ produced siderophores at an intermediate level. By contrast, siderophore production under iron-rich conditons did not differ between types and was generally low (Table S5; Figure S3). Although differences between strains from soil versus pond habitats were generally small (Figure S3), we moreover found that PVD strains from pond featured lower pyoverdine production than PVD strains from soil (t_48.1_= -2.877, p = 0.006).

**Table S4 | Growth and pyoverdine production of strain types.** Post-hoc comparisons of differences in (A) growth and (B) pyoverdine production between four strain types (PVD_PRO_, PVD, NON_PRO_ and NON). P-values are adjusted for multiple testing using the false discovery rate. Significant p-values are in bold print. Note that, independent of the comparisons below, NON strains from pond grew, on average, better than NON strains from soil (estimate ± SE: 0.09 ± 0.04, t_91.1_ = 2.252, p =0.027).

| **(A) growth** | | | | | | |
| --- | --- | --- | --- | --- | --- | --- |
| **contrast** | **condition** | **estimate** | **SE** | **df** | **t** | **p** |
| PVD_PRO_ - PVD | iron-limited | 0.09 | 0.03 | 56.23 | 2.557 | **0.016** |
| PVD_PRO_ - NON_PRO_ | iron-limited | 0.17 | 0.03 | 56.23 | 4.989 | **< 0.001** |
| PVD_PRO_ - NON | iron-limited | 0.27 | 0.03 | 56.23 | 8.017 | **< 0.001** |
| PVD - NON_PRO_ | iron-limited | 0.08 | 0.03 | 56.23 | 2.432 | **0.018** |
| PVD - NON | iron-limited | 0.18 | 0.03 | 56.23 | 5.460 | **< 0.001** |
| NON_PRO_ - NON | iron-limited | 0.10 | 0.03 | 56.23 | 3.028 | **0.006** |
| PVD_PRO_ - PVD | iron-rich | 0.04 | 0.06 | 58.22 | 0.637 | 0.632 |
| PVD_PRO_ - NON_PRO_ | iron-rich | -0.01 | 0.06 | 58.22 | -0.179 | 0.859 |
| PVD_PRO_ - NON | iron-rich | 0.11 | 0.06 | 58.22 | 1.914 | 0.182 |
| PVD - NON_PRO_ | iron-rich | -0.05 | 0.06 | 58.22 | -0.816 | 0.627 |
| PVD - NON | iron-rich | 0.07 | 0.06 | 58.22 | 1.277 | 0.413 |
| NON_PRO_ - NON | iron-rich | 0.12 | 0.06 | 58.22 | 2.093 | 0.182 |
| **(B) pyoverdine production** | | | | | | |
| **contrast** | **condition** | **estimate** | **SE** | **df** | **t** | **p** |
| PVD_PRO_ - PVD | iron-limited | 7.32 | 6.38 | 59.73 | 1.147 | 0.2558 |
| PVD_PRO_ - NON_PRO_ | iron-limited | 56.75 | 6.38 | 59.73 | 8.889 | **< 0.001** |
| PVD_PRO_ - NON | iron-limited | 71.51 | 6.38 | 59.73 | 11.2 | **< 0.001** |
| PVD - NON_PRO_ | iron-limited | 49.42 | 6.38 | 59.73 | 7.742 | **< 0.001** |
| PVD - NON | iron-limited | 64.19 | 6.38 | 59.73 | 10.05 | **< 0.001** |
| NON_PRO_ - NON | iron-limited | 14.76 | 6.38 | 59.73 | 2.313 | **0.029** |
| PVD_PRO_ - PVD | iron-rich | 1.60 | 1.88 | 58.89 | 0.85 | 0.4783 |
| PVD_PRO_ - NON_PRO_ | iron-rich | 10.40 | 1.88 | 58.89 | 5.54 | **< 0.001** |
| PVD_PRO_ - NON | iron-rich | 10.28 | 1.88 | 58.89 | 5.476 | **< 0.001** |
| PVD - NON_PRO_ | iron-rich | 8.80 | 1.88 | 58.89 | 4.69 | **< 0.001** |
| PVD - NON | iron-rich | 8.68 | 1.88 | 58.89 | 4.626 | **< 0.001** |
| NON_PRO_ - NON | iron-rich | -0.12 | 1.88 | 58.89 | -0.06 | 0.9494 |

**Table S5 | Total siderophore production of strain types.** Post-hoc comparisons of differences in total siderophore production between four strain types (PVD_PRO_, PVD, NON_PRO_ and NON). P-values are adjusted for multiple testing using the false discovery rate. Significant p-values are in bold print.

| **contrast** | **condition** | **estimate** | **SE** | **df** | **t** | **p** |
| --- | --- | --- | --- | --- | --- | --- |
| PVD_PRO_ - PVD | iron-limited | 0.11 | 0.05 | 59.05 | 2.456 | **0.020** |
| PVD_PRO_ - NON_PRO_ | iron-limited | 0.18 | 0.05 | 59.05 | 3.933 | **< 0.001** |
| PVD_PRO_ - NON | iron-limited | 0.31 | 0.05 | 59.05 | 6.596 | **< 0.001** |
| PVD - NON_PRO_ | iron-limited | 0.07 | 0.05 | 59.05 | 1.476 | 0.145 |
| PVD - NON | iron-limited | 0.19 | 0.05 | 59.05 | 4.140 | **< 0.001** |
| NON_PRO_ - NON | iron-limited | 0.12 | 0.05 | 59.05 | 2.663 | **0.015** |
| PVD_PRO_ - PVD | iron-rich | 0.01 | 0.01 | 45.81 | 0.545 | 0.588 |
| PVD_PRO_ - NON_PRO_ | iron-rich | -0.01 | 0.01 | 45.81 | -0.826 | 0.496 |
| PVD_PRO_ - NON | iron-rich | -0.02 | 0.01 | 45.81 | -1.668 | 0.307 |
| PVD - NON_PRO_ | iron-rich | -0.02 | 0.01 | 45.81 | -1.371 | 0.354 |
| PVD - NON | iron-rich | -0.03 | 0.01 | 45.81 | -2.213 | 0.192 |
| NON_PRO_ - NON | iron-rich | -0.01 | 0.01 | 45.81 | -0.842 | 0.496 |

**Table S6 | Determinants of supernatant effects.** Determinants of effects that donors had on receiver growth through compounds secreted into the supernatant under iron-limited and iron-rich conditions. Donors and receivers were isolated from soil or pond samples. Significant p-values are in bold print.

|  | **χ^2^** | **df** | **p** |
| --- | --- | --- | --- |
| condition | 419.21 | 1 | **< 0.001** |
| habitat | 3.69 | 1 | 0.055 |
| donor | 3.54 | 3 | 0.316 |
| receiver | 36.23 | 3 | **< 0.001** |
| condition : donor | 40.06 | 3 | **< 0.001** |
| condition : receiver | 6.29 | 3 | 0.098 |
| habitat : receiver | 8.57 | 3 | **0.036** |
| donor : receiver | 8.31 | 9 | 0.503 |
| condition : donor : receiver | 32.61 | 9 | **< 0.001** |

**Table S7 | Supernatant effects of strain types.** Post-hoc comparisons of differences in supernatant effects that strains of four types (PVD_PRO_, PVD, NON_PRO_ and NON) have on each other. P-values are adjusted for multiple testing using the false discovery rate. Significant p-values are in bold print. Independent of the comparisons below, PRO strains from pond overall benefitted more from receiving supernatants from others than PRO strains from soil (ratio ± SE: 1.06 ± 0.03, t_14_ = 2.233, p =0.043).

| **supernatant donor** | **supernatant receiver** | **condition** | **response** | **SE** | **t_14_** | **p** |
| --- | --- | --- | --- | --- | --- | --- |
| PVD_PRO_ | PVD_PRO_ | iron-limited | 1.72 | 0.09 | 9.784 | **< 0.001** |
| PVD | PVD_PRO_ | iron-limited | 1.38 | 0.13 | 3.341 | **0.005** |
| NON_PRO_ | PVD_PRO_ | iron-limited | 1.45 | 0.09 | 5.980 | **< 0.001** |
| NON | PVD_PRO_ | iron-limited | 1.27 | 0.07 | 4.326 | **0.001** |
| PVD_PRO_ | PVD | iron-limited | 1.46 | 0.11 | 5.152 | **< 0.001** |
| PVD | PVD | iron-limited | 1.65 | 0.08 | 10.043 | **< 0.001** |
| NON_PRO_ | PVD | iron-limited | 1.31 | 0.07 | 5.122 | **< 0.001** |
| NON | PVD | iron-limited | 1.13 | 0.05 | 2.706 | **0.017** |
| PVD_PRO_ | NON_PRO_ | iron-limited | 1.12 | 0.15 | 0.862 | 0.403 |
| PVD | NON_PRO_ | iron-limited | 1.05 | 0.09 | 0.601 | 0.557 |
| NON_PRO_ | NON_PRO_ | iron-limited | 1.50 | 0.08 | 7.392 | **< 0.001** |
| NON | NON_PRO_ | iron-limited | 1.15 | 0.05 | 3.275 | **0.006** |
| PVD_PRO_ | NON | iron-limited | 1.39 | 0.25 | 1.812 | 0.091 |
| PVD | NON | iron-limited | 1.34 | 0.19 | 2.046 | 0.060 |
| NON_PRO_ | NON | iron-limited | 1.58 | 0.20 | 3.646 | **0.003** |
| NON | NON | iron-limited | 1.49 | 0.13 | 4.490 | **0.001** |
| PVD_PRO_ | PVD_PRO_ | iron-rich | 0.98 | 0.01 | -2.693 | **0.017** |
| PVD | PVD_PRO_ | iron-rich | 0.96 | 0.02 | -2.040 | 0.061 |
| NON_PRO_ | PVD_PRO_ | iron-rich | 0.98 | 0.01 | -1.473 | 0.163 |
| NON | PVD_PRO_ | iron-rich | 1.00 | 0.01 | -0.143 | 0.889 |
| PVD_PRO_ | PVD | iron-rich | 0.98 | 0.01 | -1.725 | 0.106 |
| PVD | PVD | iron-rich | 0.99 | 0.01 | -1.303 | 0.214 |
| NON_PRO_ | PVD | iron-rich | 0.97 | 0.01 | -1.769 | 0.099 |
| NON | PVD | iron-rich | 1.00 | 0.01 | -0.038 | 0.970 |
| PVD_PRO_ | NON_PRO_ | iron-rich | 0.90 | 0.02 | -3.929 | **0.002** |
| PVD | NON_PRO_ | iron-rich | 0.92 | 0.02 | -3.978 | **0.001** |
| NON_PRO_ | NON_PRO_ | iron-rich | 0.93 | 0.01 | -5.671 | **< 0.001** |
| NON | NON_PRO_ | iron-rich | 0.93 | 0.02 | -3.449 | **0.004** |
| PVD_PRO_ | NON | iron-rich | 0.96 | 0.03 | -1.391 | 0.186 |
| PVD | NON | iron-rich | 0.93 | 0.04 | -1.613 | 0.129 |
| NON_PRO_ | NON | iron-rich | 0.97 | 0.04 | -0.885 | 0.391 |
| NON | NON | iron-rich | 0.96 | 0.03 | -1.184 | 0.256 |

**Table S8 | Determinants of community functioning.** Determinants of the variation explained in (A) community productivity and (B) pyoverdine production in series of linear models fit on data from each community (see the main text for details). P-values are adjusted for multiple testing using the false discovery rate. Significant p-values are in bold print. Note that strain identity explained more variation in productivity (t_135_ = 2.264, p = 0.025), and all predictors explained more variation in pyoverdine production (t_139_ = 6.700, p < 0.001), under iron-limited than iron-rich conditions.

|  | **(A) Productivity** | | | **(B) Pyoverdine production** | | |
| --- | --- | --- | --- | --- | --- | --- |
|  | χ^2^ | df | p | χ^2^ | df | p |
| **habitat** | 1.4 | 1 | 0.238 | 1.0 | 1 | 0.316 |
| **DOI^§^** | 133.8 | 4 | **< 0.001** | 277.8 | 4 | **< 0.001** |
| **condition** | 7.1 | 1 | **0.008** | 45.8 | 1 | **< 0.001** |
| **name : condition** | 10.4 | 4 | **0.035** | - | - | - |
| **habitat : condition** | - | - | - | 3.8 | 1 | **0.052** |
| ^§^Determinant of interest; strain identity, non-linear richness, interaction score or variable combination | | | | | | |

**Table S9 | Contributions of different strain types to pyoverdine production.** Results of linear models testing whether different strain types featured above-average or below-average contributions to community pyoverdine production under iron-limited and iron-rich conditions.

| **strain type** | **condition** | **emmean** | **SE** | **df** | **t** | **p** |
| --- | --- | --- | --- | --- | --- | --- |
| PVD_PRO_ | iron-limited | 135.90 | 19.88 | 59.94 | 6.837 | **< 0.001** |
| PVD | iron-limited | 83.84 | 19.88 | 59.94 | 4.218 | **< 0.001** |
| NON_PRO_ | iron-limited | -92.35 | 19.88 | 59.94 | -4.646 | **< 0.001** |
| NON | iron-limited | -127.38 | 19.88 | 59.94 | -6.409 | **< 0.001** |
| PVD_PRO_ | iron-rich | 45.01 | 14.35 | 59.94 | 3.136 | **0.003** |
| PVD | iron-rich | 50.37 | 14.35 | 59.94 | 3.510 | **0.001** |
| NON_PRO_ | iron-rich | -50.28 | 14.35 | 59.94 | -3.504 | **0.001** |
| NON | iron-rich | -45.10 | 14.35 | 59.94 | -3.143 | **0.003** |

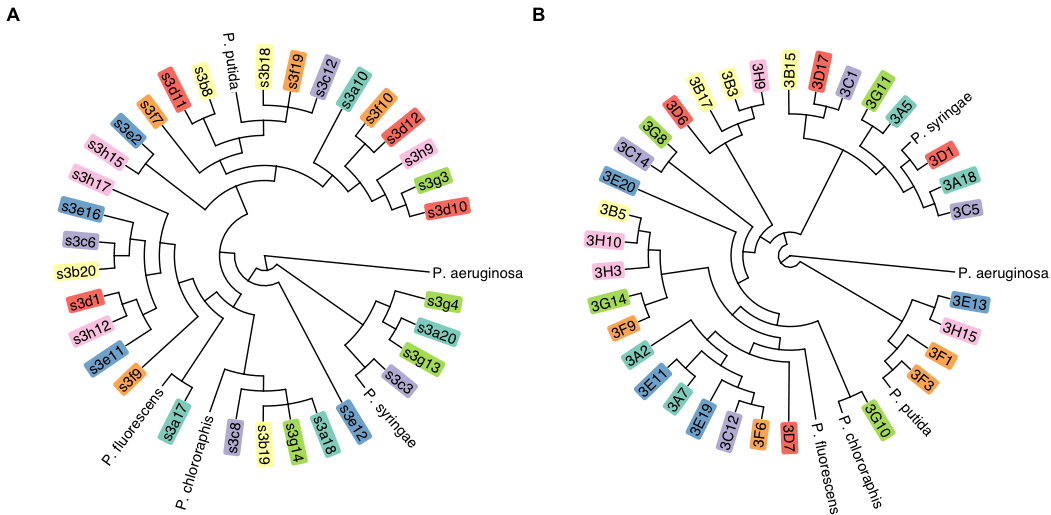

**Figure S1 | Phlyogenetic trees of natural *Pseudomonas* isolates.** Maximum-likelihood cladograms for soil (A) and pond (B) isolates based on partial rpoD sequences. For both habitats, published rpoD sequences of five well-characterised fluorescent pseudomonads were integrated into the cladograms to demonstrate taxonomic affiliation and the high diversity of our environmental isolates. Coloured rectangles represent the community from which each isolate originated.

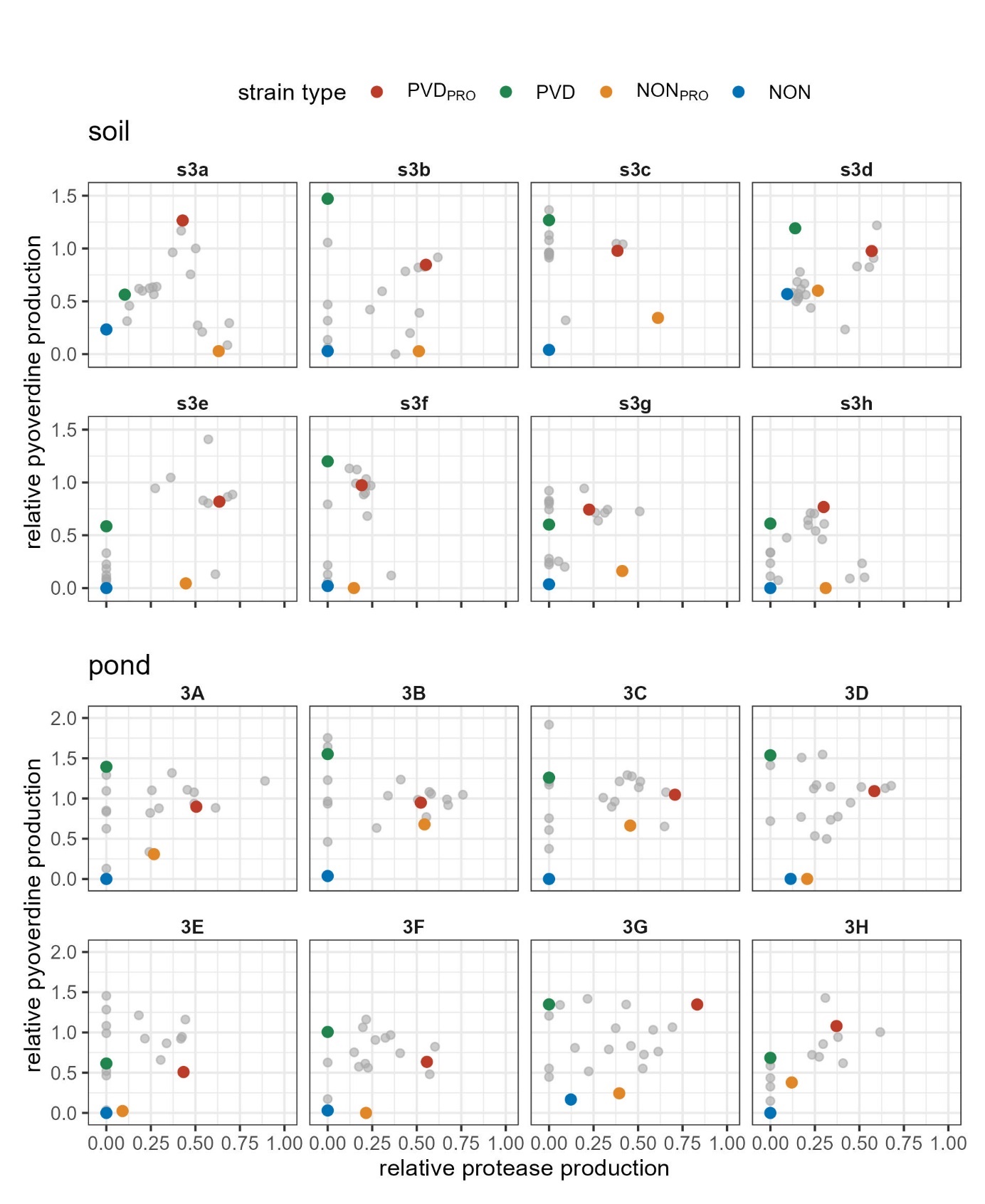

**Figure S2 | Selection of strains belonging to four different types.** We selected (i) one strain producing pyoverdine and proteases (PVD_PRO_; red), (ii) one strain producing only pyoverdine (PVD; green), (iii) one strain producing only proteases (NON_PRO_; orange), and (iv) one strain producing neither pyoverdine nor proteases (NON; blue) from each of 16 sets of 18-20 *Pseudomonas* isolates from soil and pond samples. Where ‘clean’ phenotypes were not available, we picked strains similar to the intended type. The remaining (non-selected) strains per sample (grey) are shown for comparison. Protease and pyoverdine production values are relative to those of the laboratory reference strain *P. aeruginosa* PAO1.

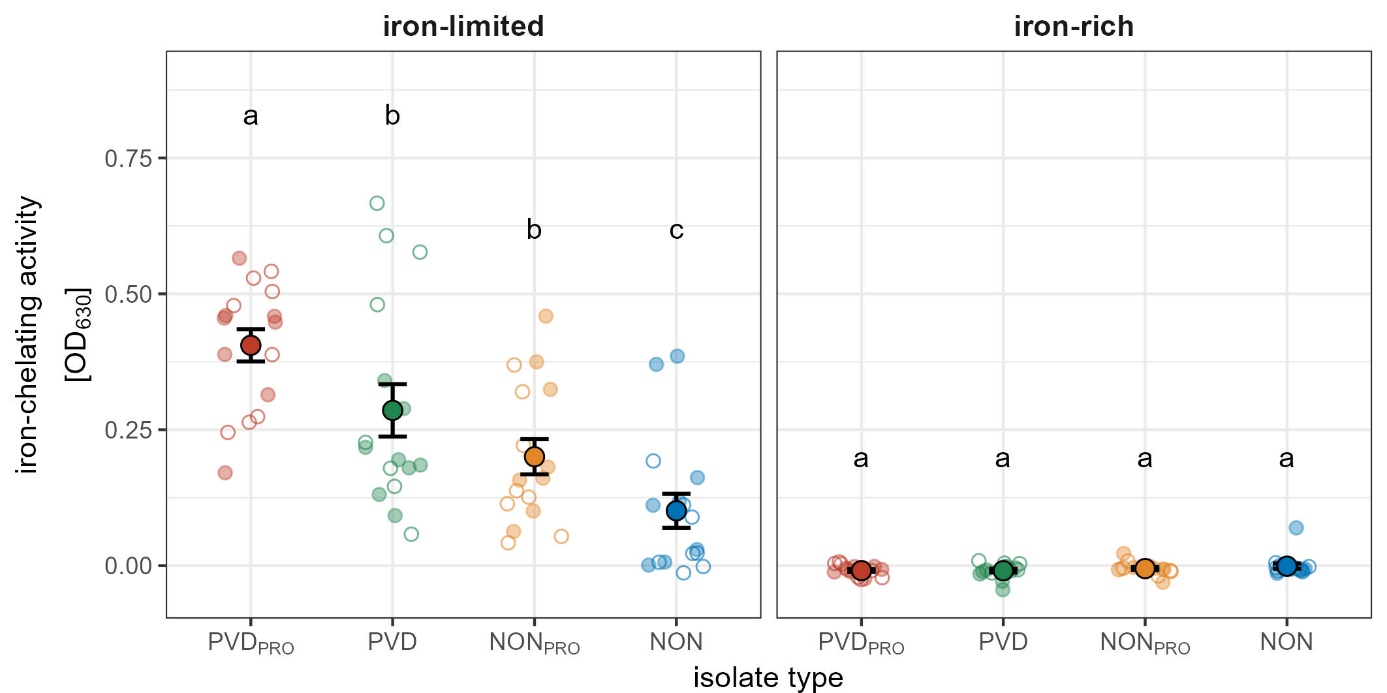
**Figure S3 | Total iron-chelating acitivity of supernatant donors.** Total iron-chelating activity of PVD_PRO_ (red), PVD (green), NON_PRO_ (orange), and NON (blue) strains isolated from eight soil (empty small circles) and eight freshwater (filled small circles) communities (one strain per type and community), measured in iron-limited and iron-rich medium. Large circles and black lines show mean and standard error, respectively. Small circles represent the the median of four replicates obtained for each strain under each condition. Letters show significantly different types. All comparisons were performed within each medium (detailed statistical results are provided in Table S3).
